# Activating the regenerative potential of Müller glia cells in a regeneration-deficient retina

**DOI:** 10.1101/180133

**Authors:** Katharina Lust, Joachim Wittbrodt

## Abstract

Regeneration responses in animals are widespread across phyla. To identify molecular players that confer regenerative capacities to non-regenerative species is of key relevance for basic research and translational approaches. Here we report a differential response in retinal regeneration between medaka (*Oryzias latipes*) and zebrafish (*Danio rerio*). In contrast to zebrafish, medaka Müller glia (olMG) cells behave like progenitors and exhibit a restricted capacity to regenerate the retina. After injury, olMG cells proliferate but fail to self-renew and ultimately only restore photoreceptors.

In our injury paradigm, we observed that in contrast to zebrafish, proliferating olMG cells do not maintain *sox2* expression. Sustained *sox2* expression in olMG cells confers regenerative responses similar to those of zebrafish MG (drMG) cells. We show that a single, cell-autonomous factor reprograms olMG cells and establishes a regeneration-like mode. Our results position medaka as an attractive model to delineate key regeneration factors with translational potential.

## Introduction

The ability to regenerate individual cells, lost organs or even the structure of the entire body is widespread in the animal kingdom. The means by which certain species achieve remarkable feats of regeneration whereas others have restricted or no capacity to do so is poorly understood. Teleost fishes are widely used models to study development, growth and regeneration of the visual system (Centanin et al., 2011; Raymond et al., 1988, 2006; Rembold et al., 2006). The retina of these fish undergoes lifelong neurogenesis, and the range of retinal cell types is generated from two sources. The first are the cells of the ciliary marginal zone (CMZ), which include retinal stem cells that give rise to progenitor cells and ultimately differentiated cell types of the growing neural retina (Centanin et al., 2011, 2014a; Raymond et al., 2006). The second source of new retinal cells are Müller glia (MG) cells, which generate new cell types during homeostasis and regeneration (Bernardos et al., 2007).

Some teleost species, including goldfish (*Carassius auratus*) and zebrafish (*Danio rerio*) have been analyzed with respect to their ability to regenerate the retina and recover visual function after injuries (Bernardos et al., 2007; Braisted and Raymond, 1992; Raymond et al., 1988; Sherpa et al., 2008). Among these, zebrafish are the most well-studied and have been shown to contain multipotent MG cells which can self-renew and regenerate all retinal neuronal and glial cell types after injuries. It is currently assumed that other teleost species posses the same regenerative capacities, however detailed analyses have been lacking.

To investigate MG cell-mediated retina regeneration in a distantly related teleost, we chose the Japanese ricefish medaka (*Oryzias latipes*), which is a well-established model organism that shared its last common ancestor with zebrafish between 200 and 300 million years ago (Schartl et al., 2013). Few regeneration studies have been carried out in medaka, but the literature reveals some interesting differences to zebrafish. Whereas fins can be fully regenerated in adult medaka (Nakatani et al., 2007), the heart has no regenerative capacity (Ito et al., 2014; Lai et al., 2017). The development and growth of the neural retina of medaka has been studied (Centanin et al., 2011, 2014b; Martinez-Morales et al., 2009), but regeneration studies are missing.

After injuries, multipotent MG cells of the zebrafish retina have been shown to upregulate the expression of pluripotency factors including *lin-28*, *oct-4*, *c-myc* and *sox2* [Ramachandran et al., 2010]. Sox2 is well known for its role in maintaining the pluripotency of embryonic stem cells (Masui et al., 2007) and is one of the four original Yamanaka factors required for the generation of induced pluripotent stem cells (Takahashi et al., 2007). Sox2 has been frequently used in reprogramming studies, such as the conversion of mouse and human fibroblasts directly into induced neural stem cells (Ring et al., 2012), or the transformation of NG2 glia into functional neurons following stab lesions in the adult mouse cerebral cortex (Heinrich et al., 2014). In the regenerating zebrafish retina, *sox2* expression is upregulated 2 days post injury (dpi) and is necessary and sufficient for the MG proliferation associated with regeneration (Ramachandran et al., 2010; Gorsuch et al., 2017).

In the present study, we find that medaka MG (olMG) cells display a restricted regenerative potential after injury and only generate photoreceptors (PRCs). We observed that olMG cells can re-enter the cell cycle after injures but fail to divide asymmetrically or generate neurogenic clusters, two steps which are essential to full regeneration. Using *in vivo* imaging, two-photon mediated specific cell ablations and lineage tracing, we find that olMG cells react preferentially to injuries of PRCs and are only able to regenerate this cell type. We demonstrate that *sox2* is expressed in MG cells in the absence of injury but, in contrast to zebrafish, is not maintained in proliferating olMG cells after injury. We show that inducing targeted expression of *sox2* in olMG cells is sufficient to shift olMG cells into a regenerative mode reminiscent of zebrafish, where they self-renew and regenerate multiple retinal cell types.

## Results

### olMG cells reenter the cell cycle after injury but do not generate neurogenic clusters

In contrast to zebrafish MG cells, olMG cells have been shown previously to be quiescent in the juvenile retina (Lust et al., 2016). In order to address the regenerative abilities of olMG cells we used the the *rx2*::H2B-eGFP transgenic line that labels the CMZ, olMG cells and cone PRCs but no rods in juvenile (8dpf) and adult medaka (Martinez-Morales et al., 2009, Reinhardt et al., 2015 and Figure S1). To investigate the reaction of olMG cells and the retina upon injury, we performed needle injuries on *rx2*::H2B-eGFP transgenic fish. To label cells re-entering the cell cycle we subsequently analyzed the fish either by immunohistochemistry for the mitotic marker phospho-histone H3 (PH3) at 3 dpi or incubated them in BrdU for 3 days to label cells in S-phase. We detected proliferating cells in the central retina, on the basis of both labels PH3 (Figures 1A-1A'') and BrdU (Figures 1B-1B'') 3 days after a needle injury. These proliferating cells were also positive for *rx2*-driven H2B-eGFP, showing that the olMG cells had re-entered the cell cycle. These results demonstrate that olMG cells in juvenile medaka are quiescent in an uninjured background (Lust et al., 2016), but begin to proliferate upon injury.

**Figure 1.**
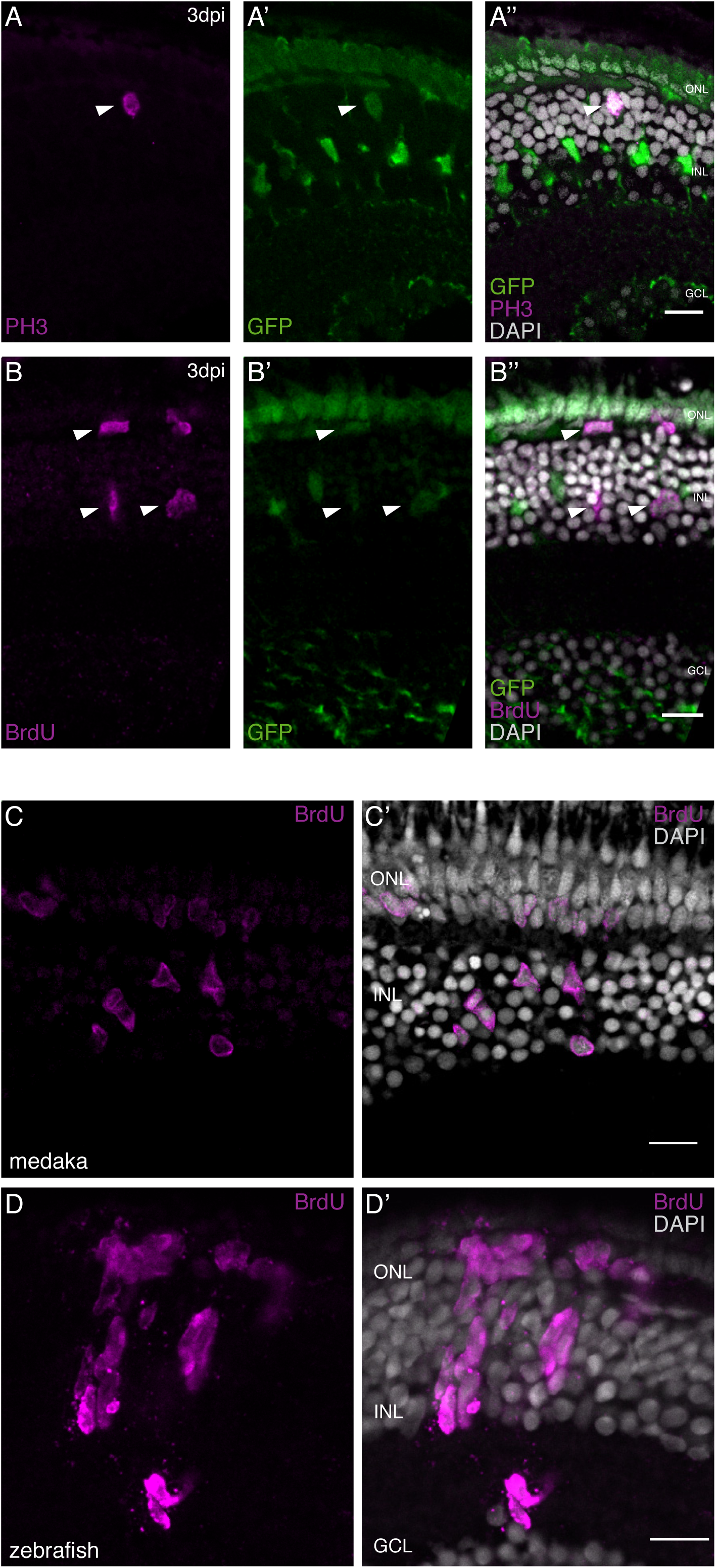
olMG cells re-enter the cell cycle after injury but do not generate neurogenic clusters. (A-A”’) Cryosections of a needle-injured juvenile medaka retina of the transgenic line *rx2*::H2B-eGFP. PH3 stainings (magenta) on the juvenile medaka retinae 3 days post needle injury show mitotic cells present in the central retina (arrowhead), co-localizing with the *rx2* nuclear reporter expression (green). (n=4 fish, data obtained from two independent experiments). (B-B”’) Cryosection of a needle-injured juvenile medaka retina of the transgenic line *rx2*::H2B-eGFP. A 3-day pulse of BrdU marks proliferating cells in the central retina after needle injury (arrowheads). BrdU staining (magenta) co-localizes with *rx2* nuclear reporter expression (green), indicating that olMG cells re-entered the cell cycle. (n=6 fish, data obtained from three independent experiments). (C, C’) Cryosection of a needle-injured juvenile medaka retina. BrdU-positive (magenta) single cells are present in the INL and ONL. (n=6 fish, data obtained from two independent experiments). (D, D’) Cryosection of a needle-injured zebrafish retina. BrdU-positive (magenta) neurogenic clusters are present in the INL. Additionally, BrdU-positive proliferating cells can be detected in the ONL (n=3 fish, data obtained from two independent experiments). Scale bars are 10 μm.

The onset of MG proliferation in zebrafish has been observed between 1 and 2 dpi (Fausett and Goldman, 2006). To understand if olMG cells show a similar mode of activation, we performed BrdU incorporation experiments and analyzed time-points after injury ranging from 1 dpi until 3 dpi. At 1 dpi, no BrdU-positive cells were detected in the retina (data not shown). At 2 dpi, the first BrdU positive cells were detected in the INL and the outer nuclear layer (ONL) of the central retina (Figure S2A-S2B‴). Co-localization with GFP showed that these cells are olMG cells or olMG-derived cells (Figure S2A'' and S2B'').

In response to injury olMG cells initiate DNA synthesis and divide maximally once as indicated by the appearance of single or a maximum of two BrdU-positive cells next to each other in the INL at both 2 dpi and 3 dpi (Figures 1C and 1C’).

In contrast the injury response of zebrafish MG (drMG) cells at comparable juvenile stages (4dpf) is characterized by the formation of large nuclear, neurogenic clusters in the INL (Figures 1D and 1D’). This is consistent with the response of adult drMGs to injury in which a single asymmetric division produces a MG cell and a progenitor cell that divides rapidly to generate neurogenic clusters (Nagashima et al., 2013).

These results show that olMG cells start re-entering the cell cycle between 1 and 2 dpi but do not generate neurogenic clusters.

### olMG cells react preferentially to PRC injuries by apical migration

For proper regeneration to occur, the appropriate cell types must be produced. This requires not only the regulation of the proliferation of stem or progenitor cells, but also the proper control of lineage decisions in the progenitors. If and when fate decisions are made by the MG cells or proliferating progenitors during regeneration is largely unknown. To study whether different injury sites (PRC or retinal ganglion cell (RGC) injury) result in a differential response of olMG cells, we used two-photon mediated ablations and consecutive imaging (Figures S3A-S3D) and addressed their behavior in immediate (up to 30 hpi) and late (until 6dpi) response to injury.

We induced PRC injuries in medaka and observed that MG nuclei below the wound site started migrating apically at 17 hours post injury (hpi) (Figures 2A-2A”’, see also Movie S1). These migrations were not coordinated between individual cells. Some nuclei migrated into the ONL, whereas others stayed at the apical part of the INL. Nuclei farther from the wound site did not migrate in response to the injuries. In contrast, after RGC injuries, there was no migration of MG nuclei, either apically or basally toward the wound, within the first 30 hpi (Figures 2B-2B”’, see also Movie S1).

**Figure 2.**
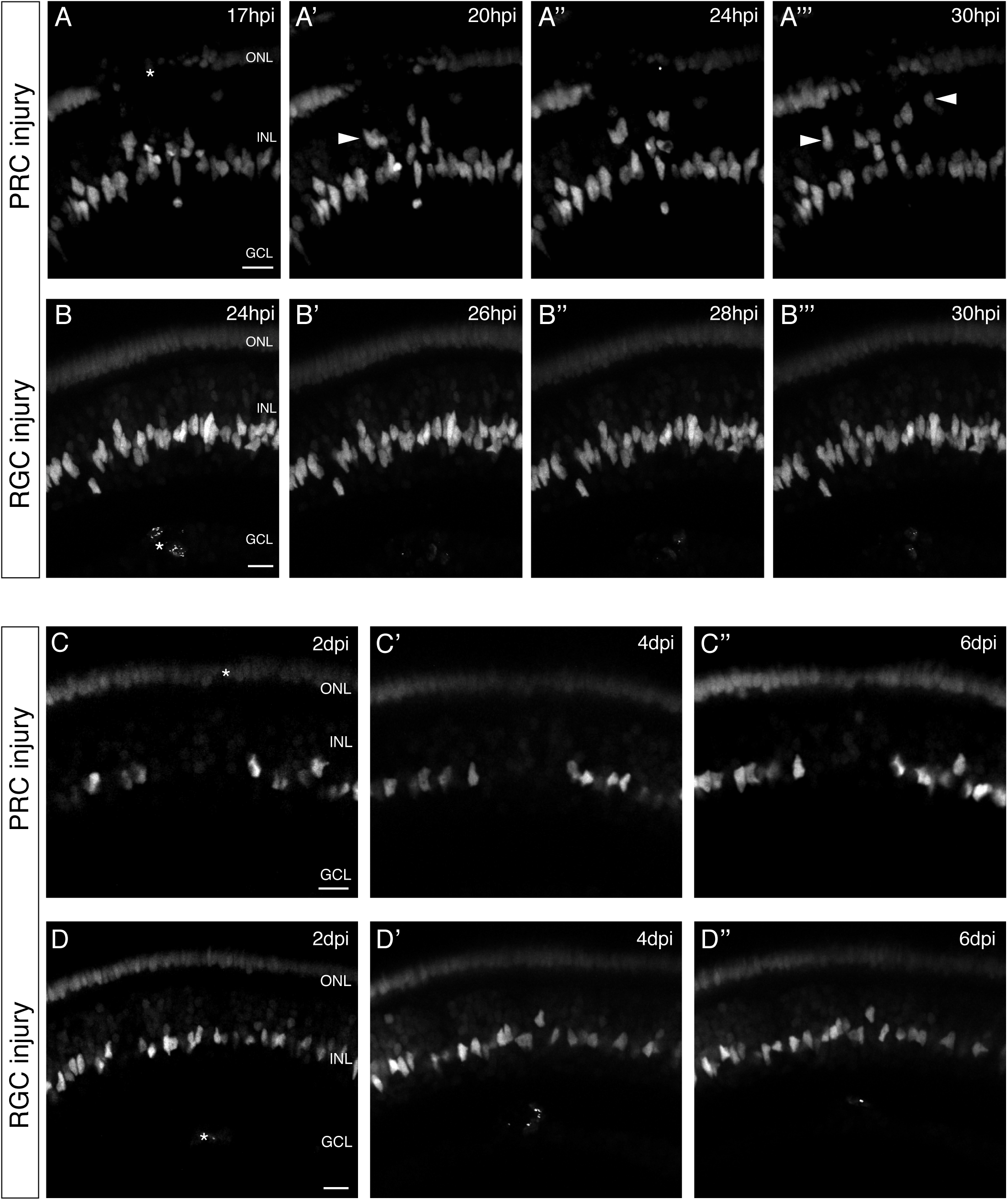
olMG cells react preferentially to PRC injuries by apical migration. (A-B”’) *In vivo* imaging of juvenile *rx2*::H2B-eGFP medaka retinae which were either injured in the ONL or the GCL (asterisks) using a two-photon laser and imaged consecutively until 30 hpi (n>10 fish each, data obtained from >10 independent experiments each). (A-A”’) After PRC injuries olMG nuclei (arrowheads) start migrating apically towards the ONL layer from 17 hpi on. The migration is not coordinated among different migrating nuclei. (B-B”’) After RGC injuries no migration of olMG nuclei can be detected until 30 hpi. Scale bars are 10 μm. (C-D’) *In vivo* imaging of juvenile *rx2*::H2B-eGFP medaka retinae which were either injured in the ONL or the GCL (asterisks) using a two-photon laser and imaged every second day after injury (n>10 fish each, data obtained from >10 independent experiments each). (C-C’) PRC injuries result in an apical migration of olMG nuclei into the injury site. The following days until 6 dpi the nuclei do not migrate back towards the INL resulting in a gap of MG nuclei in the INL. (D-D’) After RGC injuries no migration of olMG nuclei can be detected until 6 dpi. Scale bars are 10 μm.

To investigate whether medaka MG nuclei migrate back at later time-points after PRC injuries or show any migratory behavior after RGC injuries, we re-imaged the injury site at two-day intervals to follow an injured retina up to 6 dpi. At 2 dpi, retinae with PRC injuries showed a gap in the INL below the injury site, at a position where MG nuclei are normally found, reflecting the migration MG nuclei towards the ONL from this location (Figures 2C-4C’). The gap in the INL persisted until 6 dpi (Figure 2C’). The reaction of olMG cells in retinae with RGC injuries differed. Here, we neither observed an apical nor basal migration of olMG nuclei (Figures 2D-2D’) and in fact no migration of olMG nuclei was observed at all until 6 dpi. To rule out that this is due to too little damage in the RGC layer we increased the injury size. This lead to swelling and secondary cell death in PRCs and activated olMG cells to migrate apically (Figure S4A-S4B), indicating further that their preferential reaction is towards PRC injuries.

Taken together, these results show that olMG cell nuclei migrate towards PRC injury sites within 24 hpi and remain in this location up until 6 days, whereas they display no discernible reaction towards RGC injuries. This indicates a clear preferential reaction of olMG nuclei to refill the injured PRC layer.

### olMG nuclei but not their cell bodies are depleted after PRC injuries

Long-term *in vivo* imaging of fish that were injured in the ONL made it apparent that olMG nuclei migrate apically into the wound site but remain there which might indicate a complete remodeling of the soma of these neuroepithelial cells. To understand whether cell bodies of the olMG cells remain intact during this nuclear migration, we observed nuclear movements (transgenic line *rx2*::H2B-eGFP) in the context of the olMG cell body (transgenic line *rx2*::lifeact-eGFP). We imaged the animals at two-day intervals following ONL injuries. As previously observed, olMG nuclei migrated out of the INL into the wound site (Figures 3A-3A’). Cell bodies of the olMG cells spanning the entire apico-basal distance remained intact until 6 dpi in the absence of a nucleus in the INL (Figure 3A’). The previous site of the nucleus could still be detected by a small enlargement of the soma in this region.

**Figure 3.**
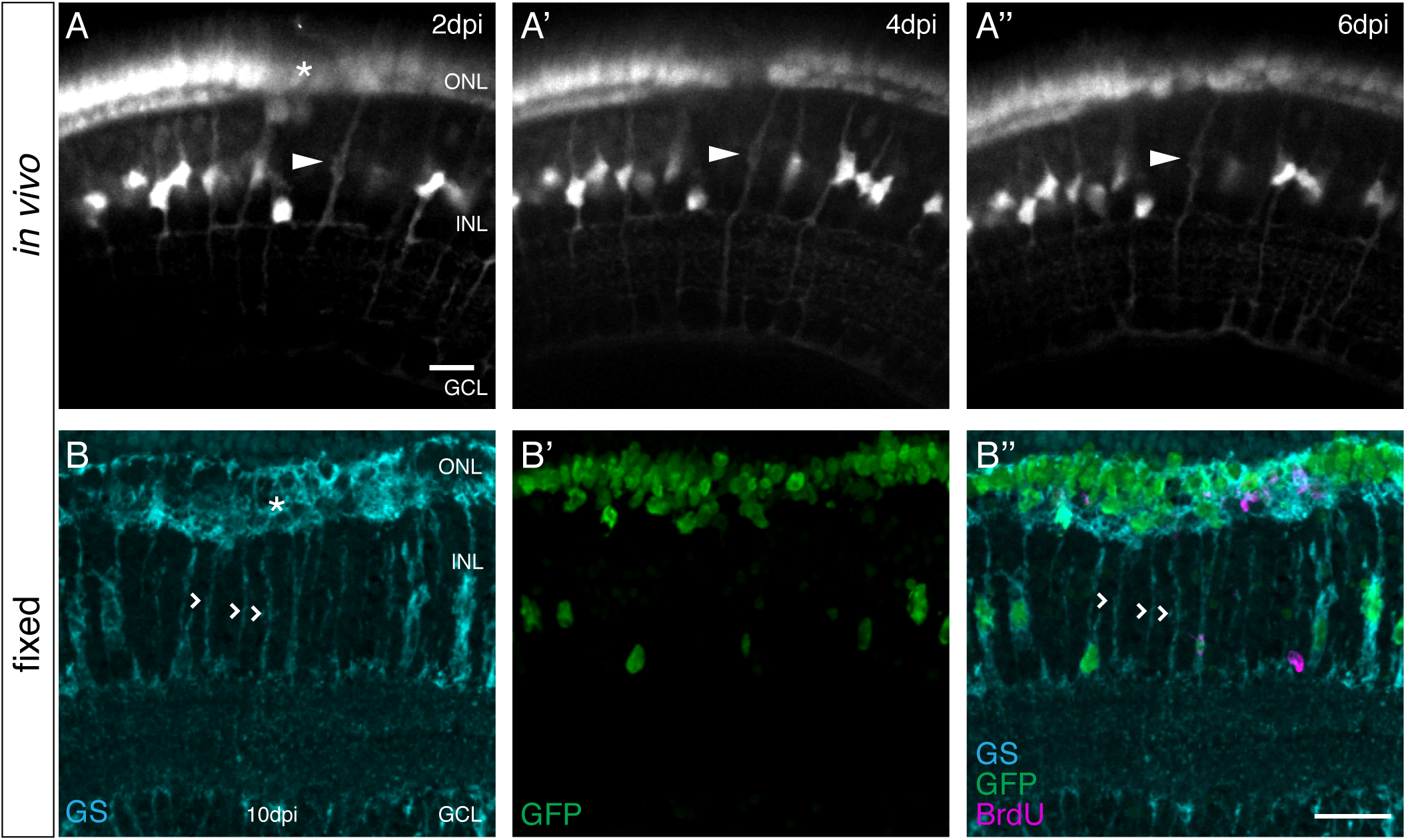
olMG nuclei are depleted after PRC injuries without process loss. (A-A’) *In vivo* imaging of a juvenile *rx2*::H2B-eGFP, *rx2*::lifeact-GFP medaka retina which was injured in the ONL (asterisk) and imaged every second day after injury. Close to the injury site an olMG process without a nucleus can be detected at 2 dpi (A, arrowhead). The empty process remains until 6 dpi (A”) (n=3 fish, data obtained from three independent experiments). Scale bar is 10 μm. (B-B’) Cryosections of the injured juvenile medaka retina of the transgenic line *rx2*::H2BeGFP. The fish were injured in the ONL (asterisk), incubated in BrdU for 3 days and fixed at 10dpi. Many GFP-positive nuclei (green) are located in the ONL, some co-localizing with BrdU (magenta). In the INL few GFP-positive nuclei are present. Many GS-positive (turquoise) olMG processes below the injury site do not contain a GFP-positive nucleus. Next to the empty processes GFP-positive nuclei can be detected within GS-positive processes (n=4 fish, data obtained from two independent experiments). Scale bar is 20 μm.

Additionally, to extend the time range of analysis, we performed immunohistochemistry on fish injured in the ONL. After incubation in BrdU for 3 days and fixation at 10 dpi, we observed similar results (Figures 3B-3B’). In this procedure, we used immunohistochemistry to detect GS to label the MG processes (Figure 3B). BrdU-positive cells in the ONL mark the location of the injury (Figure 3B’). In the region directly below, the majority of MG nuclei, which had been labelled by *rx2*::H2B-eGFP, were absent from the INL (Figure 3B’). GS-positive cell bodies remained spanning the apico-basal width, but without the apparent presence of nuclei. In contrast, unaffected GS-positive olMG cells located on either side of the wound site still contained their nuclei, as could easily be detected by the large size of the soma. This data shows that the cell bodies of injury-activated olMG cells are still intact despite the migration of their nuclei into the ONL.

### olMG cells divide in the INL with an apico-basal distribution

Since the injury response of olMG cells apparently does not involve self-renewal of olMG cells we wondered about the position and orientation of the cell division plane, a factor which has been associated with cell fate in various systems.

We first addressed the apico-basal position of dividing olMG cells by PH3 immunohistochemistry after injury. We detected PH3-positive cells only in the INL (Figures 4A-4A’). Some dividing cells were located more apically (Figures 4A-4A’), while others were located more basally (data not shown). This is in contrast to findings in zebrafish where, in a light injury paradigm, PH3-positive MG cells can be found in the ONL 2 days after injury (Nagashima et al., 2013)

**Figure 4.**
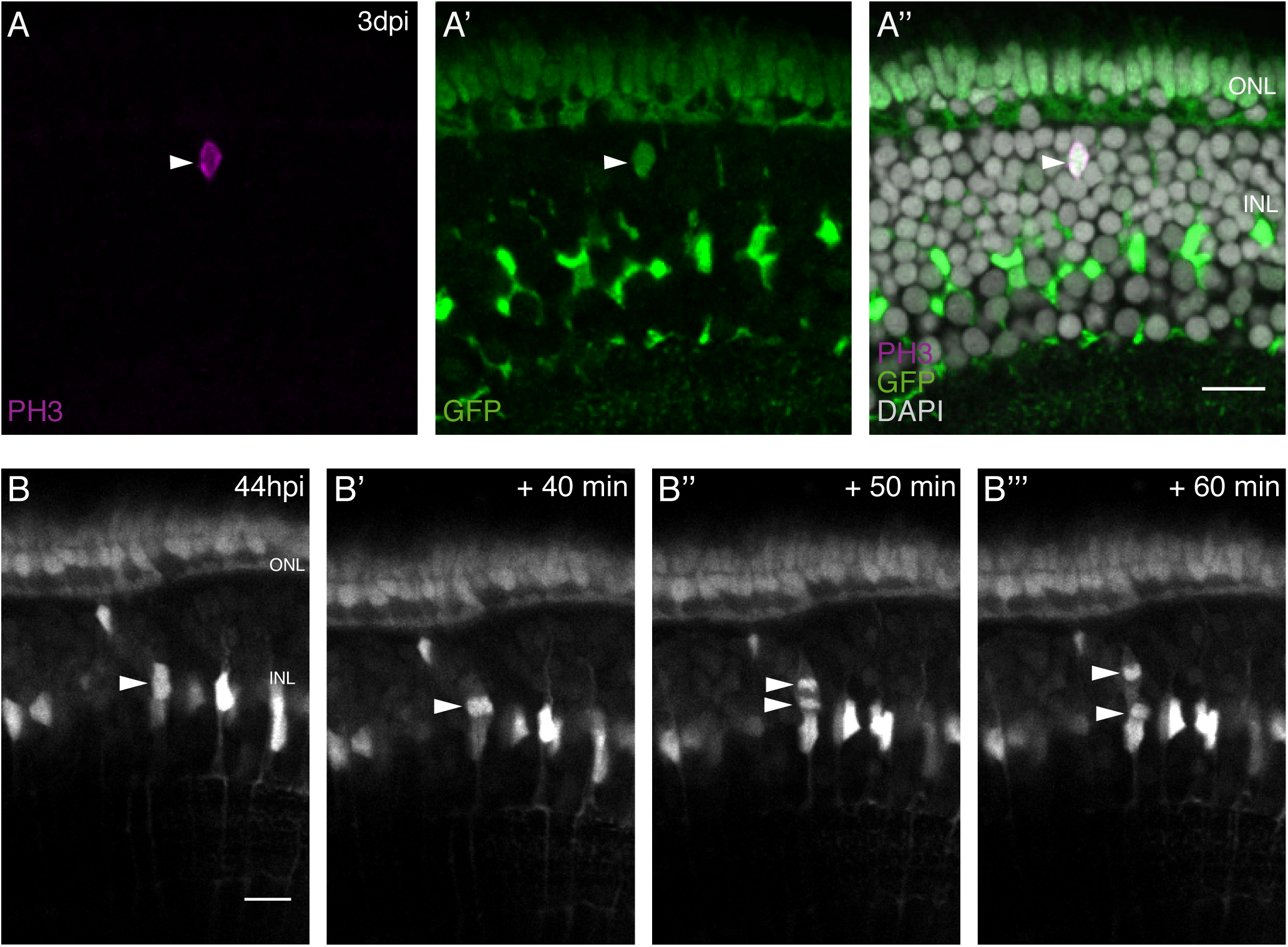
olMG cells divide in the INL with an apico-basal spindle orientation. (A-A’) Cryosection of an injured juvenile medaka retina of the transgenic line *rx2*::H2B-eGFP. PH3 stainings (magenta) on juvenile medaka retinae 3 days post PRC injury show mitotic olMG cells present in the INL (arrowhead), co-localizing with the *rx2* nuclear reporter expression (green) (n=4 fish, data obtained from three independent experiments). Scale bars are 10 μm. (B-B”’) *In vivo* imaging of juvenile *rx2*::H2B-eGFP medaka retinae which were injured in the ONL and imaged starting at 44 hpi. OlMG nuclei which start to condense their chromatin can be detected in the INL (arrowheads). The divisions occur in an apico-basal manner (n=6 fish, data obtained from six independent experiments, 5 out of 6 imaged divisions were apico-basal). Scale bars are 10 μm.

To address the cleavage plane of dividing olMG cells we employed *in vivo* imaging of *rx2*::H2BeGFP fish, which permits visualizing the separation of chromatin and thus gives a measurement of the orientation of division. The first injury-triggered olMG divisions were observed at 44 hpi (Figures 4B-4B‴, see also Movie S3). They occurred in the INL, both in the center and close to the ONL (data not shown). The mode of division was preferentially apico-basal (5 out of 6 cases), while only a single horizontal division was observed (1 out of 6 cases). These results show that injury induced olMG cell divisions occur in different positions in the INL and have a strong preference to occur apico-basally.

### olMG cells are lineage restricted

In zebrafish drMG cells are able to regenerate all neuronal cell types and self-renew after injury (Nagashima et al., 2013; Powell et al., 2016). We followed a BrdU-based lineage tracing approach successfully applied in zebrafish (Fausett and Goldman, 2006; Powell et al., 2016) to address the potency of olMG cells. Transgenic *rx2*::H2B-eGFP fish retinae were injured either by two-photon laser ablation of PRCs or RGCs specifically or using a needle ablating all cell types. The injured fish were incubated in BrdU for 3 days to label proliferating cells. This allows to efficiently detect all injury triggered S-phase entry of olMG cells (Figure S5A-S5D). For lineaging, fish were grown until 14 dpi to allow a regeneration response and subsequently analyzed for BrdU-positive cells in the different retinal layers (Figure 5A). PRC injuries led to the detection of 97% of all BrdU-positive cells in the ONL, mostly in the rod nuclear layer, indicative for PRC fate (Figures 5B and 5E). No BrdU-positive cells could be detected in the INL. Additionally, we found that the INL below the injury site was devoid of olMG cell nuclei, both consistently arguing for the absence of injury triggered olMG self-renewal. Strikingly, RGC injuries did not trigger BrdU-uptake in olMG cells or any other differentiated cell type (data not shown). Needle injuries affecting all retinal cell types triggered the same response as the specific lesions in the PRC layer. 97% of all BrdU-positive cells were present in the ONL, and only a single BrdU-positive olMG cell was found in 1 of 10 fish (Figures 5C and 5E). Also later application of BrdU after injury (4 to 7 dpi) did not result in BrdU-positive MG cells (Figure S6A-S6C). Importantly, BrdU-positive nuclei were not positive for GS, indicating that they were not MG cells anymore (Figure 5D), but were positive for Recoverin, a PRC marker (Figure 5E). These results demonstrate that olMG cells do not self-renew and rather function as mono-potent repair system restricted to the generation of PRCs, most of which belong to the rod lineage.

**Figure 5.**
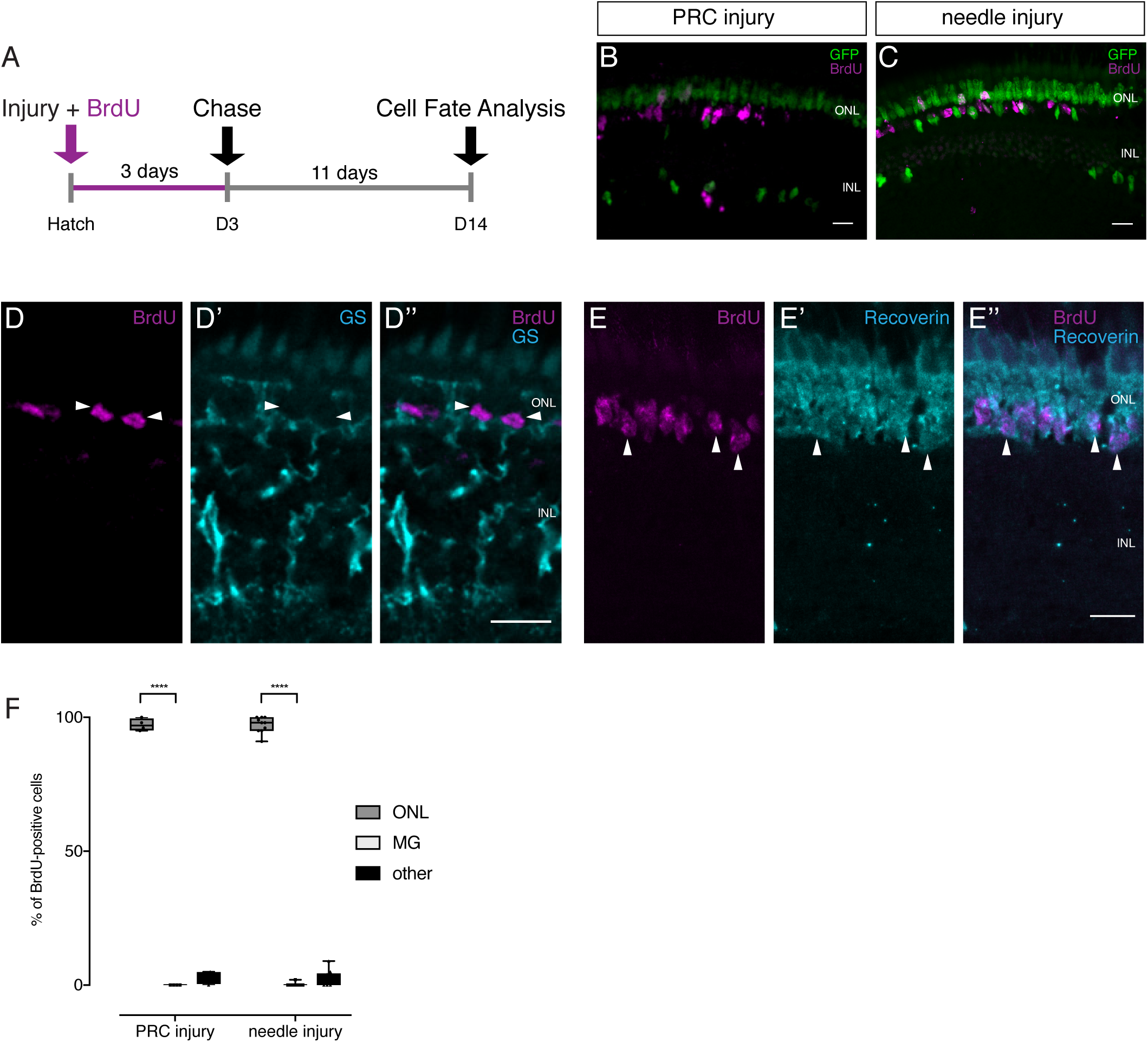
Lineage tracing after injuries reveals the preferential regeneration of PRCs. (A) Scheme outlining the experimental procedure. Juvenile medaka were injured in the retina with a two-photon laser ablating either PRCs or RGCs or with a needle ablating all cell types. The fish were incubated in BrdU for 3 days and analysed at 14 dpi. (B-C) PRC injuries result in BrdU-positive cells in the ONL, mostly in the rod layer. No BrdU-positive olMG cells are present and fewer GFP-positive MG cells are found in the INL (n=4 fish, data obtained from two independent experiments). Needle injuries result in BrdU-positive cells in the ONL, mostly in the rod layer. Except for 1 BrdU-positive olMG cell in 1 fish, no BrdU-positive olMG cells are detected. GFP-positive olMG nuclei are largely depleted from the INL (n=10 fish, data obtained from three independent experiments). (D-D’) After needle injuries BrdU-positive cells (magenta) in the ONL are not co-labelled with GS (cyan), indicating that they are not MG cells (n=8 fish, data obtained from two independent experiments). (E-E’) After needle injuries BrdU-positive cells (magenta) in the ONL are co-labelled with Recoverin (cyan), indicating that they PRCs (n=5 fish, data obtained from one experiments). Scale bars are 10 μm. (F) Quantification of the location of BrdU-positive reveals that in all injury types BrdU-positive cells are predominantly located in the ONL. ****p<0.0001. Box plots: median, 25th and 75th percentiles; whiskers show maximum and minimum data points.

### *Sox2* expression is not maintained in proliferating olMG cells after injury

The previous results show that olMG cells re-enter the cell cycle after injuries introduced by needle to the complete retina or by 2-photon ablation to the PRC layer. They regenerate PRC but do not undergo self-renewal. This suggests that olMG cells lack intrinsic factors that trigger self-renewal and multi-potency upon injury. One transcription factor which is well known for its involvement in the self-renewal of stem cells – particularly neural stem cells – is Sox2 (Sarkar and Hochedlinger, 2013). It has been shown that cells expressing *sox2* are capable of both self-renewal and the production of a range of differentiated neuronal cell types (Sarkar and Hochedlinger, 2013). Data from zebrafish have shown that a ubiquitous gain of Sox2 expression triggers a regenerative response of the drMGs in the absence of injury (Gorsuch et al., 2017).

To investigate the expression of *sox2* in MG cells, we performed immunohistochemistry on uninjured retinae in medaka and zebrafish. In the medaka retina, Sox2 protein is detected the in the CMZ (data not shown) as well as in amacrine cells (ACs) and olMG cells (Figure 6A-6A‴). In zebrafish, the pattern was similar: Sox2 protein was present in the CMZ (data not shown), in ACs and drMG cells (Figure 6B-6B‴). This data is consistent with data from other vertebrates including human, whose MG cells also maintain *sox2* expression (Gallina et al., 2014).

**Figure 6.**
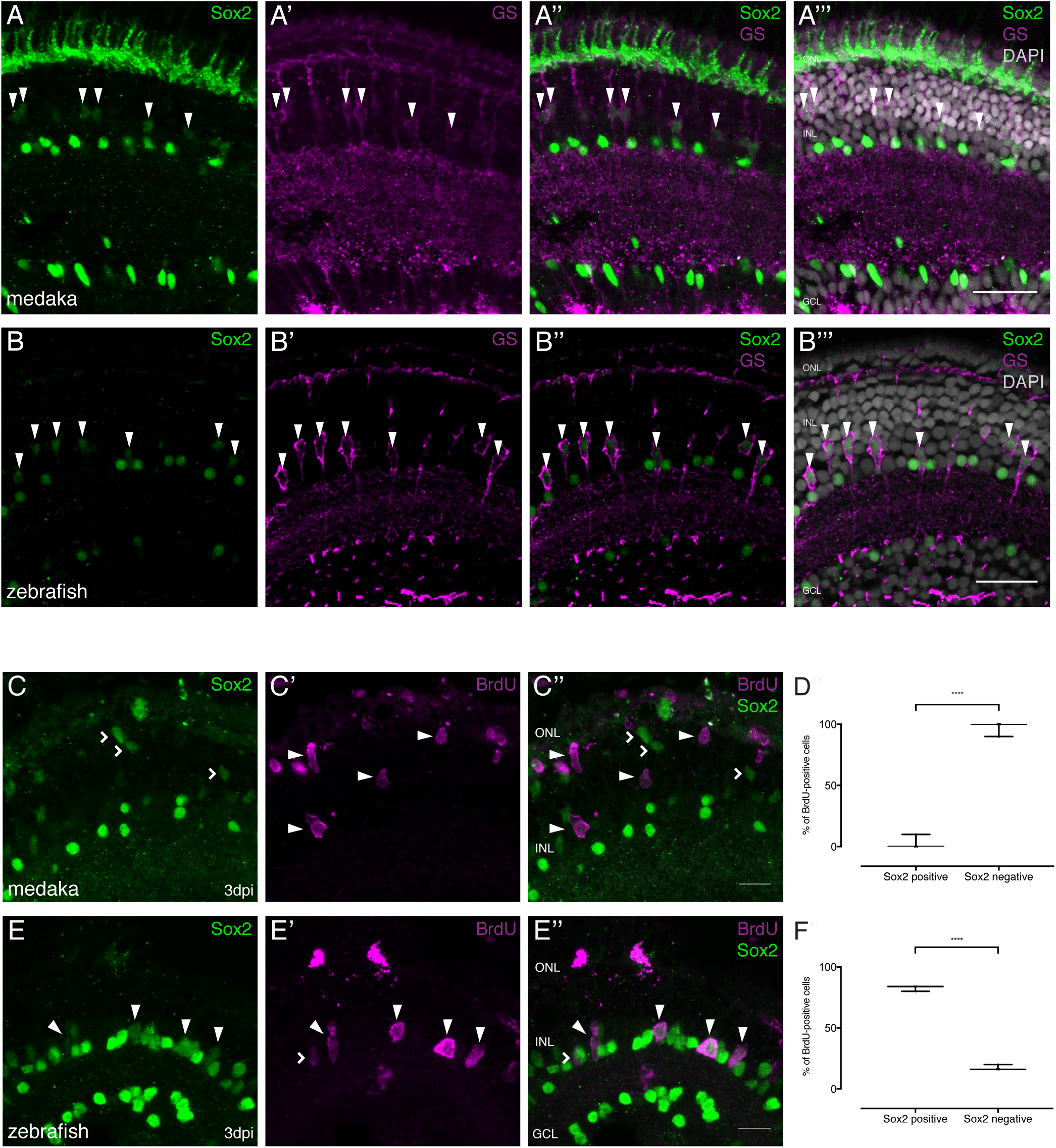
Sox2 is present in MG cells of the juvenile medaka and zebrafish retina but not maintained after injury in medaka. (A-A”’) Cryosection of an uninjured juvenile medaka retina. Sox2 (green) labeled cells with round nuclei are present in the INL and the GCL. Sox2-labeled cells with round nuclei are ACs present in the INL and the GCL. Sox2-positive cells with elongated nuclei are present in the INL (arrowheads). Co-labeling with GS (magenta) proves that cells with elongated nuclei are olMG cells. Additional staining, which is likely unspecific staining since *sox2* mRNA cannot be detected there (Reinhardt et al., 2015), can be detected in the ONL. (B-B”’) Cryosections of an uninjured zebrafish retina at 9 dpf. Sox2 (green) labeled cells with round nuclei are present in the INL and the GCL. Sox2-positive cells with elongated nuclei are present in the INL (arrowheads). Co-labeling with GS (magenta) proves that cells with elongated nuclei are drMG cells. Scale bars are 20 μm. (C-C’) Cryosection of an injured juvenile medaka retina at 3 dpi. BrdU (magenta, arrowheads) labeled cells are not co-labeled with Sox2 (green, arrowheads). Sox2-positive cells with elongated nuclei, indicating non-proliferative olMG cells, are found in the INL (open arrowheads) (n=3 fish, data obtained from two independent experiments). (D) Quantification of the amount of Sox2-positive and negative proliferating cells of BrdU-positive cells at 3 dpi in medaka. ****p<0.0001. Box plots: median, 25th and 75th percentiles; whiskers show maximum and minimum data points. (E-E’) Cryosection of an injured zebrafish retina at 3 dpi. BrdU (magenta) and Sox2 (green) double positive cells can be detected in the INL (arrowheads). BrdU-positive Sox2-negative cells can rarely be detected (open arrowhead) (n=3 fish, data obtained from two independent experiments). Scale bars are 10 μm. (F) Quantification of the amount of Sox positive and negative proliferating cells of BrdU-positive cells at 3 dpi in zebrafish. ****p<0.0001. Box plots: median, 25th and 75th percentiles; whiskers show maximum and minimum data points.

To investigate the expression of *sox2* after injuries in olMG and drMG cells respectively, we performed needle injuries, incubated the fish in BrdU and fixed them between 1 and 4 dpi. Proliferating olMG cells exhibited a downregulation in the expression of *sox2*, which was mostly absent by 3 dpi (Figures 6C-6D, 6% of all BrdU positive cells were Sox2-positive). Conversely, in zebrafish, *sox2* expression could still be detected after 3 days in drMG cells that strongly proliferated in response to injury (Figures 6E-6F, 84% of all BrdU positive cells were Sox2-positive). These findings strongly argue that the downregulation of Sox2 expression restricts the regenerative properties of olMG cells.

### Sustained Sox2 expression restores olMG driven regeneration

The results presented above indicate that after injury, olMG cells and MG-derived progenitors do not maintain an expression of *sox2,* in contrast to the situation in zebrafish. We hypothesize that the prolonged *sox2* expression facilitates drMG cells to undergo self-renewal and to generate neurogenic clusters and ultimately all cell-types necessary to regenerate a functional retina.

To test this hypothesis, we chose the inducible LexPR system (Emelyanov and Parinov, 2008) targeted to olMG cells to sustain *sox2* expression. In those retinae, we ablated all retinal cell types by a needle injury and performed BrdU mediated lineage tracing as described above. To induce the persistent expression of *sox2* in olMG cells after injuries, we employed the LexPR transactivation system (*rx2*::LexPR OP::*sox2*, OP::H2B-eGFP) allowing to follow individual *sox2* expressing cells by the nuclear eGFP expression. We induced *sox2* expression for two days and provided BrdU in parallel, performed a needle injury to ablate all cell types and maintained the expression of *sox2* until 3 dpi. After a chase until 14 dpi the retinae and regenerated cell types were analyzed (Figures 7A and 7B). In needle injured wildtype fish, olMG cells did not self-renew and only gave rise to PRCs (Figure 5). In contrast, olMG cells experiencing persistent expression of sox2 showed self-renewal and differentiation into different cell types in the ONL and INL as indicated by BrdU lineage tracing. In particular, the olMG cells maintaining Sox2 expression after the injury regenerated olMG cells, exhibited an increase in regenerated cone PRC and also formed rod PRCs and ACs (Figures 7B-7C’).

**Figure 7.**
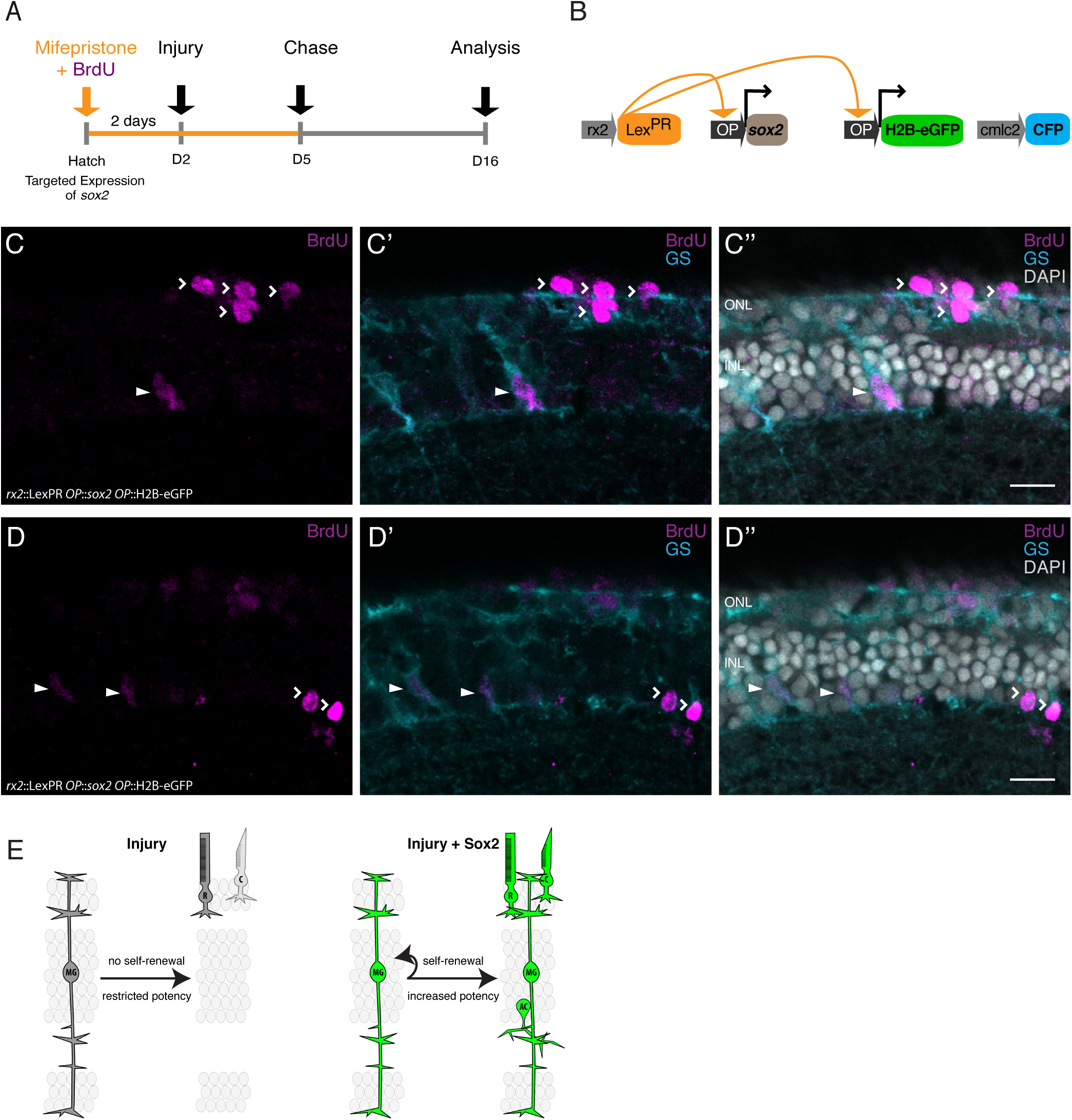
Sox2 induces a regeneration response in olMG cells. (A-B) Induction scheme and construct (B) used for *sox2* induction. (C-D’) Cryosection of an induced juvenile medaka retina. BrdU-positive (magenta) olMG cells, which are labelled by GS (cyan) can be detected in the INL (arrowheads). Additional BrdU-positive cells are located in the ONL, in the location of both rods and cones (open arrowheads), as well as the INL, in the location of ACs (open arrowheads) (n= 2 fish, data obtained from two independent experiments). Scale bars are 10 μm. (E) OlMG cells respond to injuries by proliferation without self-renewal and restriction towards PRC fate. Targeted expression of *sox2* induces self-renewal and increased potency to olMG cells.

These data indicate that a targeted maintenance of *sox2* expression after injury is sufficient to induce self-renewal and increase potency in MG cells in the medaka retina turning a mono-potent repair system into a regeneration system with increased potency.

## Discussion

Here, we have characterized a differential regenerative response between two teleost fish and used it as a framework to address the molecular determinants of regeneration during evolution. By using a combination of *in vivo* imaging, targeted cell type ablation and lineage tracing, we investigated the dynamics of the injury response in the medaka retina. We focused on MG cells, which play a prominent role in zebrafish retinal regeneration. While upon injury olMG cells re-enter the cell cycle, they fail to undergo self-renewal. Furthermore, olMG cells do not generate the neurogenic clusters which arise in zebrafish, nor do they produce all neuronal cell types in the retina. We traced this effect prominently to Sox2, the expression of which is maintained in proliferating drMG cells after injury, but not in olMG cells. We demonstrated that the sustained expression of *sox2* is sufficient to convert an olMG into a dr-like MG. The fact that this response is acquired cell autonomously and in the context of a non-regenerative retina can be relevant for putative translational approaches.

Since olMG cells did not self-renew after injuries and only had the capacity to regenerate PRC, olMG cells are not true multipotent retinal stem cells. Instead, olMG cells should be considered lineage-restricted progenitors. They re-entered the cell cycle between 1 and 2 dpi, similar to the re-entry observed in zebrafish. This indicates that the signals that are essential for cell-cycle re-entry are present in medaka and are activated in a window of time similar to that of zebrafish. However, since olMG cells did not self-renew, the first cell division is presumably symmetric. Given extensive data that the orientation of the division axis is associated with the developmental potency of the dividing cell (Cayouette and Raff, 2003; Das and Storey, 2012), the apico-basal division axis of olMG cells in response to injury provides further evidence for the non-self-renewing progenitor nature of olMG cells. Strikingly, both the developing and the regenerating zebrafish retina exhibit nuclear divisions axes of either progenitors or drMG with predominant lateral orientation, correlating with the asymmetric outcome of the division (Das et al., 2003; Lahne et al., 2015). The fact that olMG cells divided apico-basally after injuries and regenerated only PRCs indicates that their division plane (perpendicular to division plane in drMG cells) correlates with a symmetric division into two differentiating cells, depleting the olMG cell pool. This depletion could be detrimental for the retina, since MG cell bodies play a crucial role in mechanical stability of the retina (MacDonald et al., 2015) as well as light guiding through the retina (Franze et al., 2007). After retinal injures olMG cell bodies were maintained in the absence of a nucleus in the INL reflecting the necessity to preserve this structural/optical element.

In the uninjured retina, olMG cells express *sox2*, as is the case for many other vertebrates, including humans. However, *sox2* expression in olMG cells is downregulated in response to injury, in contrast to the injury response of drMG cells, which upregulate *sox2* (Gorsuch et al., 2017). We speculate that this upregulation is due to epigenetic modifications of the *sox2* locus. A recent study in the mouse retina showed that the expression of *oct4* is upregulated shortly after injury and then downregulated at 24 hpi (Reyes-Aguirre and Lamas, 2016). This correlates with a decrease in the expression of DNA methyltransferase 3b and its subsequent upregulation at 24 hpi, triggering a decrease in methylation and subsequent re-methylation of *oct4*. Furthermore, a recent study on zebrafish regeneration discovered the existence of so-called tissue regeneration enhancer elements (TREEs) (Kang et al., 2016). One TREE was associated with leptin b, which is expressed in response to injuries of the fin and heart. This TREE acquires open chromatin marks after injury, can be divided into tissue-specific modules and can drive injury-dependent expression in mouse tissue. This raises the possibility that the *sox2* locus in olMG cells experiences epigenetic modifications after injury which differ from modifications in zebrafish. The fact that *sox2* expression is detected in all vertebrate MG cells analyzed to date in the absence of injury raises the question whether a decrease in *sox2* expression after injury might be a common feature of non-regenerative species, like chicken, mouse and even humans. Data from a conditional *sox2* knockout in mouse shows that Sox2 is necessary for maintenance of MG morphology and quiescence (Surzenko et al., 2013). While its expression is maintained in response to the injection of growth factors after retinal damage (Karl et al., 2008) its regulation in response to injury alone has not been described. Data obtained in cultures of human MG cells (Bhatia et al., 2011) provide additional important insights. Strikingly similar to medaka, silencing the expression of *sox2* caused MG cells to lose stem and progenitor cell markers and adopt a neural phenotype (Bhatia et al., 2011). These findings align well with the results from medaka presented here und suggest that olMG cells and their behavior as progenitor cells can serve as a model for mammalian and in particular human MG cells.

The results shown here may provoke an evolutionary question: is retinal regeneration an ancestral or derived feature within the infraclass of teleosts? The question might be resolved by investigations of this capacity in other fish species more closely related to medaka, such as *Xiphophorus maculatus*, whose last common ancestor with medaka lived around 120 million years ago (Schartl et al., 2013). Additionally, species like the spotted gar, whose lineage diverged from teleosts before teleost genome duplication (Braasch et al., 2016), might provide insights about the ancestral mode of retinal regeneration. Recently, the retinal architecture of the spotted gar has been analyzed (Sukeena et al., 2016). Here, proliferative cells have been detected in the central retina likely representing proliferating MG cells, which generate rod PRCs during homeostasis as seen in zebrafish, suggesting that regeneration is indeed an ancestral feature in the ray-finned fish lineage.

With a potential translational perspective, regenerating and non-regenerating systems can now be systematically compared to delineate the underlying factors and mechanisms. To date, our cumulative results show that the regenerative potential of olMG cells in the context of homeostasis and injury in medaka resemble that of mammals and birds more than zebrafish. We propose that this provides an added value to medaka as a model species for regeneration studies that bridge the differences between zebrafish and mammals. Studies of heart regeneration that have compared zebrafish and medaka lend additional support this statement (Ito et al., 2014; Lai et al., 2017). As re-programmable multipotent retinal stem cells, MG cells harbor a great potential for treating degenerative retinal diseases. Our work indicates that the addition of a single re-programming factor facilitates a regeneration-like response mediated by olMG cells. Their multiple resemblances of features of mammalian and human MG cells position them as an ideal model for the development of new treatments preventing the degeneration and initiating the regeneration of the retina.

## Author Contributions

K.L. and J.W. conceived the study and designed the experiments. K.L. performed all the experiments. K.L. and J.W. wrote the manuscript.

## Acknowledgements

We thank the Wittbrodt department for constructive discussions on the project; L. Centanin and A. Gutierrez-Triana for valuable input on the project and the manuscript; N. Aghaallaei, C. Becker, F. Caroti, A.-K. Heilig, I. Krämer, S. Lemke, C. Lischik, T. Tavhelidse and E. Tsingos for critical reading of the manuscript; R. Hodge for manuscript editing. We are grateful to A. Saraceno, E. Leist and M. Majewski for fish husbandry. K.L. is a member of HBIGS, the Heidelberg Graduate School for Life Sciences and was supported by a LGFG Fellowship. This work was supported by the European Research Council (GA 294354-ManISteC to J.W.)

## Methods

### Animals and transgenic lines

Medaka (*Oryzias latipes*) and zebrafish (*Danio rerio*) used in this study were kept as closed stocks in accordance to Tierschutzgesetz 111, Abs. 1, Nr. 1 and with European Union animal welfare guidelines. Fish were maintained in a constant recirculating system at 28°C on a 14 h light/10 h dark cycle (Tierschutzgesetz 111, Abs. 1, Nr. 1, Haltungserlaubnis AZ35–9185.64 and AZ35–9185.64/BH KIT). The following stocks and transgenic lines were used: wild-type Cabs, *rx2*::H2B-eGFP, *rx2*::lifeact-GFP, *rx2*::H2B-eGFP QuiH, *rx2*::LexPR *OP*::*sox2 OP*::H2B-eGFP *cmlc2*::CFP, AB/AB zebrafish and Albino zebrafish. All transgenic lines were created by microinjection with Meganuclease (I-SceI) in medaka embryos at the one-cell stage, as previously described (Thermes et al., 2002).

### BrdU incorporation

For BrdU incorporation, fish were incubated in 2.5 mM BrdU diluted in 1x Embryo Rearing Medium (ERM) or 1x Zebrafish Medium for respective amounts of time.

### Induction of the Lex^PR^ system

For induction of the Lex^PR^ system, fish were induced by bathing them in a 5 μM to 10 μM Mifepristone solution in Embryo Rearing Medium (ERM) for respective times.

### *In vivo* imaging and laser ablations

For *in vivo* imaging fish in a Cab background were kept in 5x 1-phenyl-2-thiourea (PTU, Sigma) in 1x ERM from 1 dpf until imaging to block pigmentation. Fish in a QuiH background could be imaged without any treatment. Fish were anesthetized in 1x Tricaine diluted in 1xERM and mounted in glass bottomed Petri dishes (MaTek) in 1% Low Melting Agarose. The specimens were oriented lateral, facing down, so that the right eye was touching the cover-slip at the bottom of the dish. Imaging and laser ablations were performed on a Leica SP5 equipped with a Spectra Physics Mai Tai^®^ HP DeepSee Ti:Sapphire laser, tunable from 690-1040nm and Leica Hybrid Detectors. A wound was introduced using the bleach point function or the region of interest function, together with the high energy 2-photon laser tuned to 880nm. Follow-up imaging was performed using same laser at 880nm and a 40x objective.

### Retinal needle injuries

Larvae (zebrafish 5dpf, medaka 8dpf) were anesthetized in 1x Tricaine (A5040, Sigma-Aldrich) in 1x ERM and placed on a wet tissue. Under microscopic visualization, the right retina was stabbed multiple times in the dorsal part with a glass needle (0.1 mm diameter). Left retinae were used as controls.

### Immunohistochemistry on cryosections

Fish were euthanized using Tricaine and fixed over night in 4% PFA, 1xPTW at 4°C. After fixation samples were washed with 1x PTW and cryoprotected in 30% sucrose in 1xPTW. To improve section quality, the sections were incubated in a half/half mixture of 30% sucrose and Tissue Freezing Medium for at least 3 days. 16 μM thick serial sections were obtained on a cryostat. Sections were rehydrated in 1x PTW for 30 min at room temperature. Blocking was performed for 1-2 h with 10% NGS (normal goat serum) in 1xPTW at room temperature. The respective primary antibodies were applied diluted in 1% NGS o/n at 4°C. The secondary antibody was applied in 1% NGS together with DAPI (1:500 dilution in 1xPTW of 5mg/ml stock) for 2-3 h at 37°C. Slides were mounted with 60% glycerol and kept at 4°C until imaging.

### Antibodies

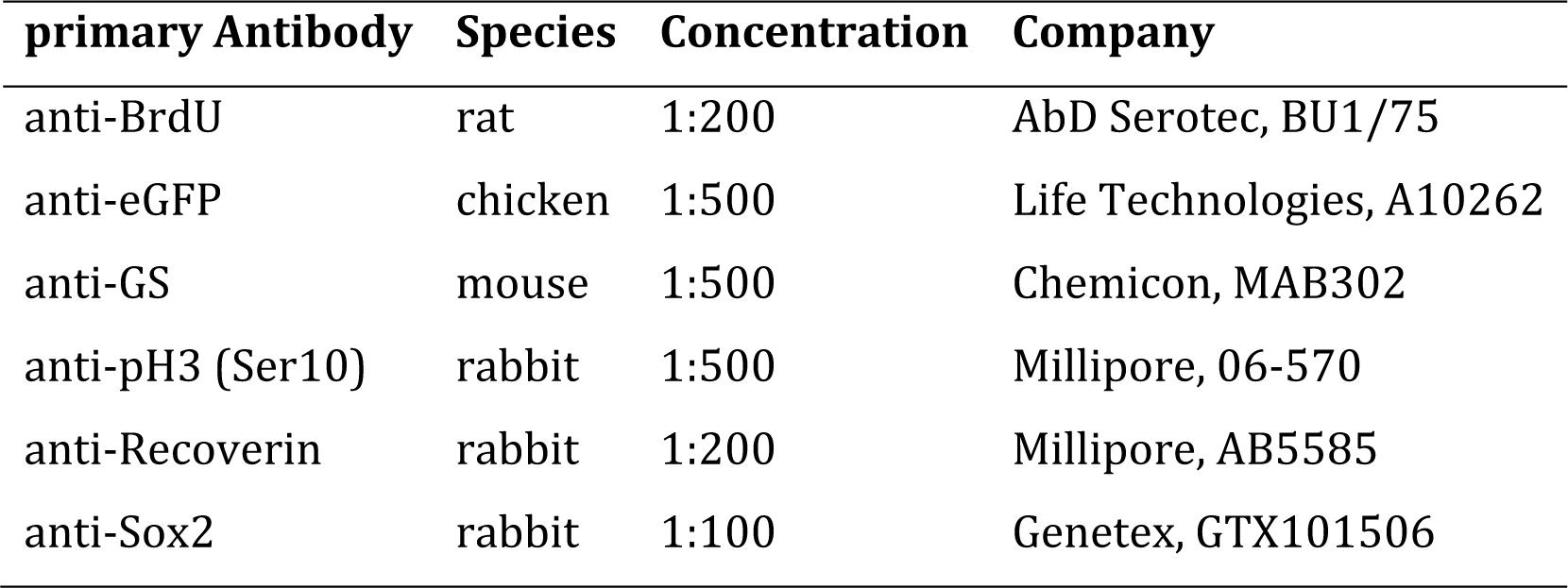

| **secondary Antibody** | **Species** | **Concentration** | **Company** |
| --- | --- | --- | --- |
| anti-chicken DyLight488 | donkey | 1:750 | Jackson, 703-485-155 |
| anti-mouse Alexa546 | goat | 1:750 | Life Technologies, A-11030 |
| anti-mouse Cy5 | donkey | 1:750 | Jackson, 715-175-151 |
| anti-rabbit DyLight549 | goat | 1:750 | Jackson, 112-505-144 |
| anti-rabbit 647 | goat | 1:750 | Life Technologies, A-21245 |
| anti-rat DyLight549 | goat | 1:750 | Jackson, 112-505-143 |
| anti-rat Alexa633 | goat | 1:750 | Life Technologies, A-21094 |

### BrdU immunohistochemistry

BrdU antibody staining was performed with an antigen retrieval step. After all antibody stainings, except for BrdU and DAPI staining, were complete a fixation for 30 min was performed with 4% PFA. Slides were incubated for 1 h at 37°C in 2 N HCl solution, and pH was recovered by washing with a 40% Borax solution before incubation with the primary BrdU antibody.

### TUNEL staining

TUNEL stainings on cryosections were performed after all other antibody stainings were completed using the In Situ Cell Death Detection Kit TMR Red by Roche. Stainings were performed according to the manufacturers protocol with the following modifications. Washes were performed with 1x PTW instead of PBS.

### Immunohistochemistry imaging

All immunohistochemistry images were acquired by confocal microscopy at a Leica TCS SPE with either a 20x water objective or a 40x oil objective.

### Image processing and statistical analysis

Images were processed via Fiji image processing software. Statistical analysis and graphical representation of the data were performed using the Prism software package (GraphPad).

### Multimedia Files

Movie S1, related to Figure 2. *In vivo* imaging of olMG nuclei reactions to a PRC injury

Movie S2, related to Figure 2. *In vivo* imaging of olMG nuclei reactions to a RGC injury

Movie S3, related to Figure 4. *In vivo* imaging of an olMG division after PRC injury

## Supplemental Figures

**Figure S1, related to Figure 1.**
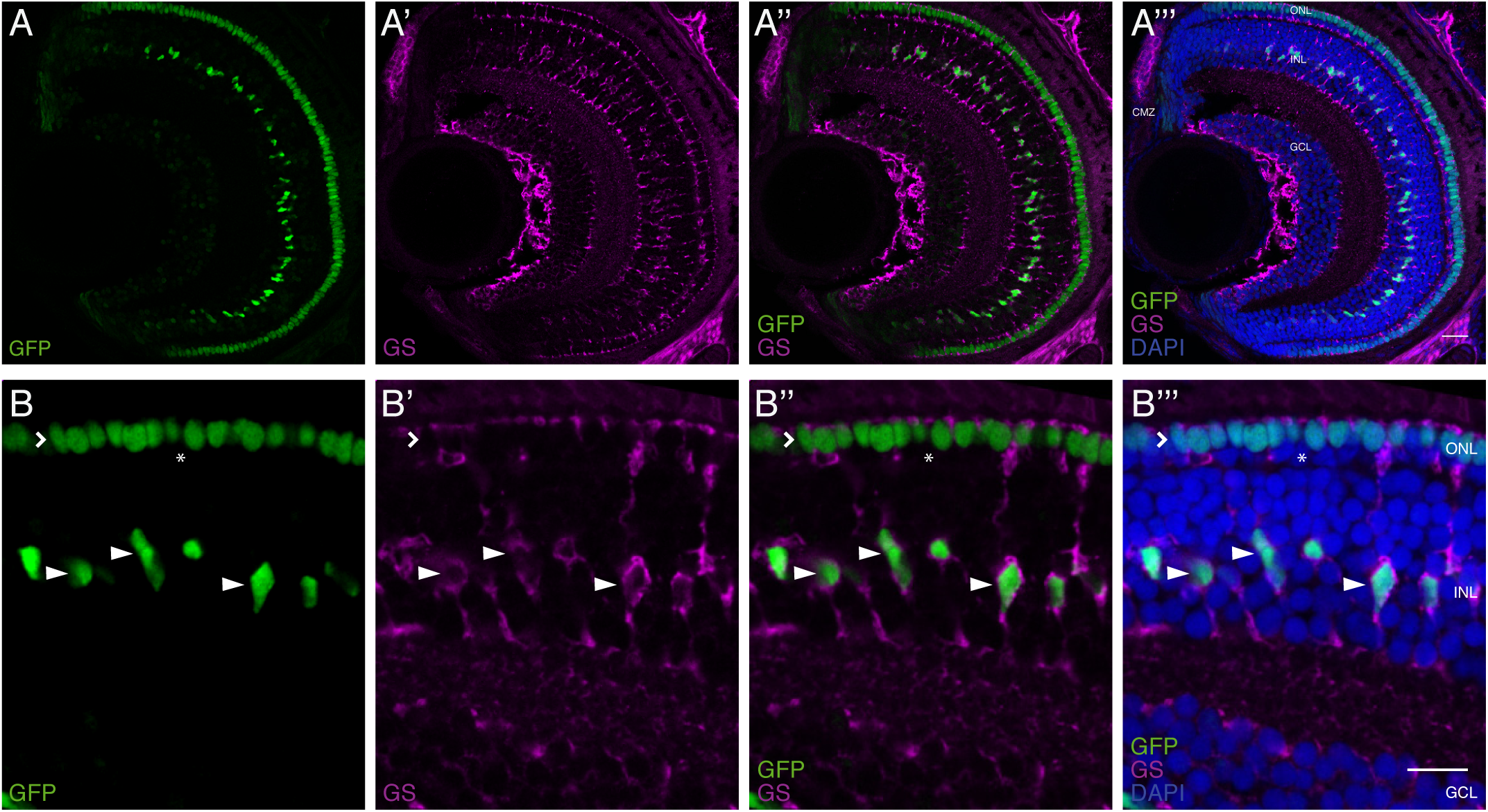
*Rx2*-reporter labels olMG cells, cone PRCs and CMZ cells in the juvenile medaka retina. (A-B”’) Cryosections of a juvenile medaka fish of the transgenic line *rx2*::H2B-eGFP. GFP-positive nuclei (green) are located in the INL, the ONL and the CMZ. GFP-positive nuclei in the INL overlap with Glutamine Synthetase (GS)-labeling (magenta, arrowheads), indicating that these cells are olMG cells. GFP-positive nuclei in the ONL are only present in the outer most nuclear layer (open arrowhead), indicating that these are cone PRCs. The inner layer of the ONL, where rod PRCs are located, is not labelled (asterisk). Scale bars are 10 μm.

**Figure S2, related to Figure 1.**
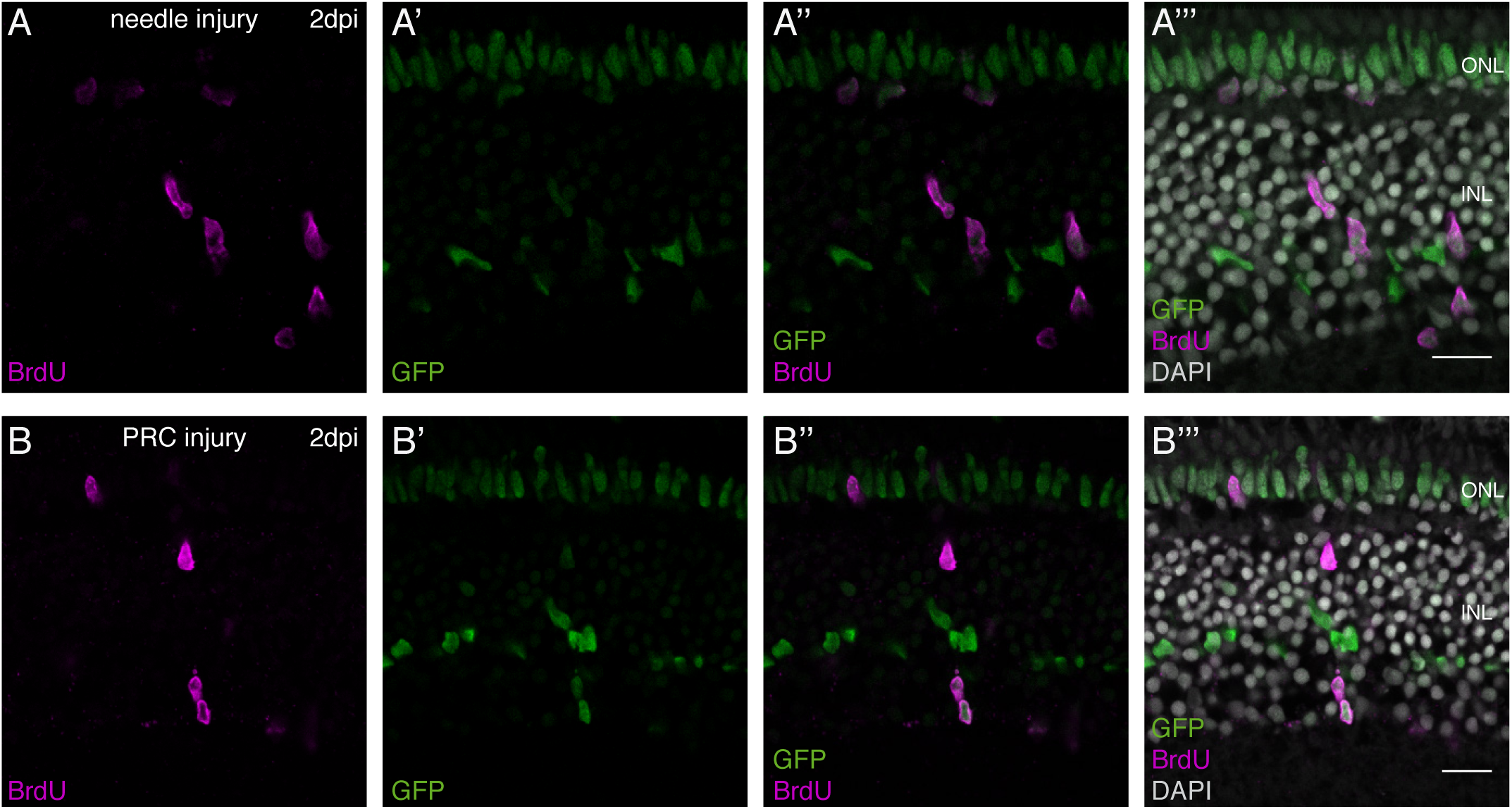
Injury-induced timing of olMG cell cycle re-entry. (A-B”’) Cryosections of either a needle-injured juvenile medaka retina (A-A”’) or a PRC-injured juvenile medaka retina (B-B”’) of the transgenic line *rx2*::H2B-eGFP. At 2 dpi, the first BrdU-positive (magenta) cells are detected in the central retina. BrdU co-localizes with *rx2*-driven GFP (green) in the INL and ONL (n=3 fish each, data obtained from two independent experiments each). Scale bars are 10 μm.

**Figure S3, related to Figure 2.**
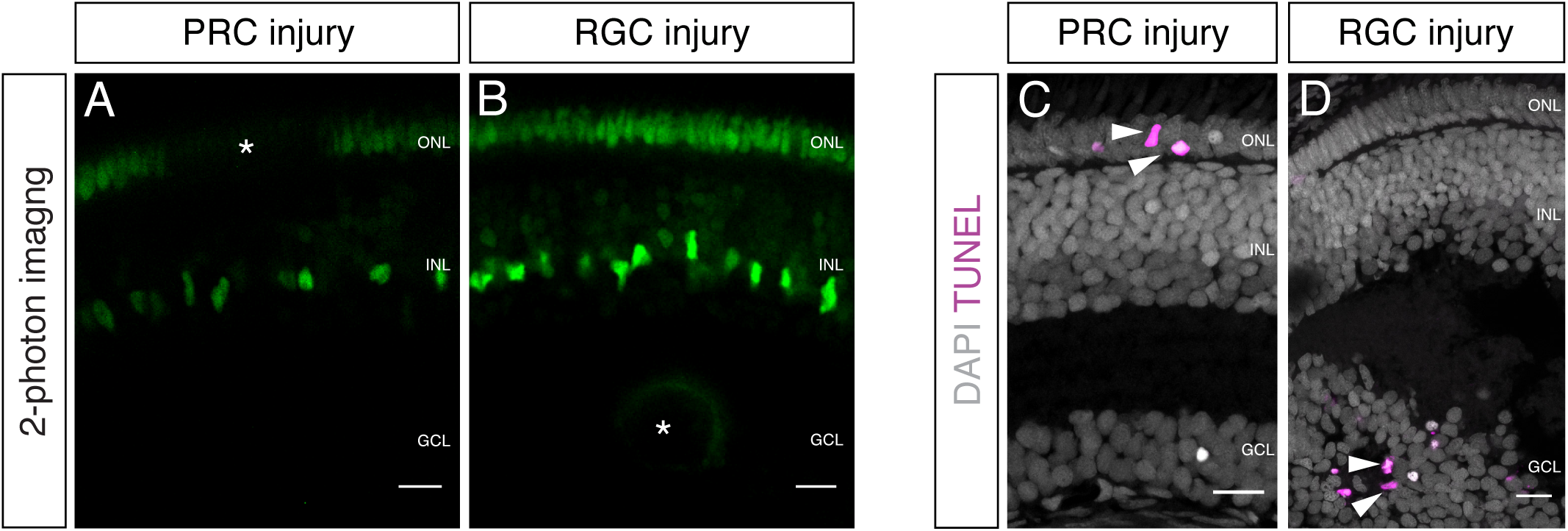
Two-photon mediated laser ablation enables targeted cell ablation in the retina resulting in specific cell death signatures. (A-B) *In vivo* imaging of juvenile *rx2*::H2B-eGFP medaka retinae which were either injured in the ONL or the GCL (asterisks) using a two-photon laser. Targeted cell type ablation can be achieved; PRCs (A) as well as cells of the GCL (B) can be ablated. (C-D) Cryosections of juvenile medaka retinae which were either injured in the ONL or the GCL and fixed 16 hpi. TUNEL stainings (magenta) to detect programmed cell death show specific cell death of either ONL (C) or GCL (D). Scale bars are 10 μm.

**Figure S4, related to Figure 2.**
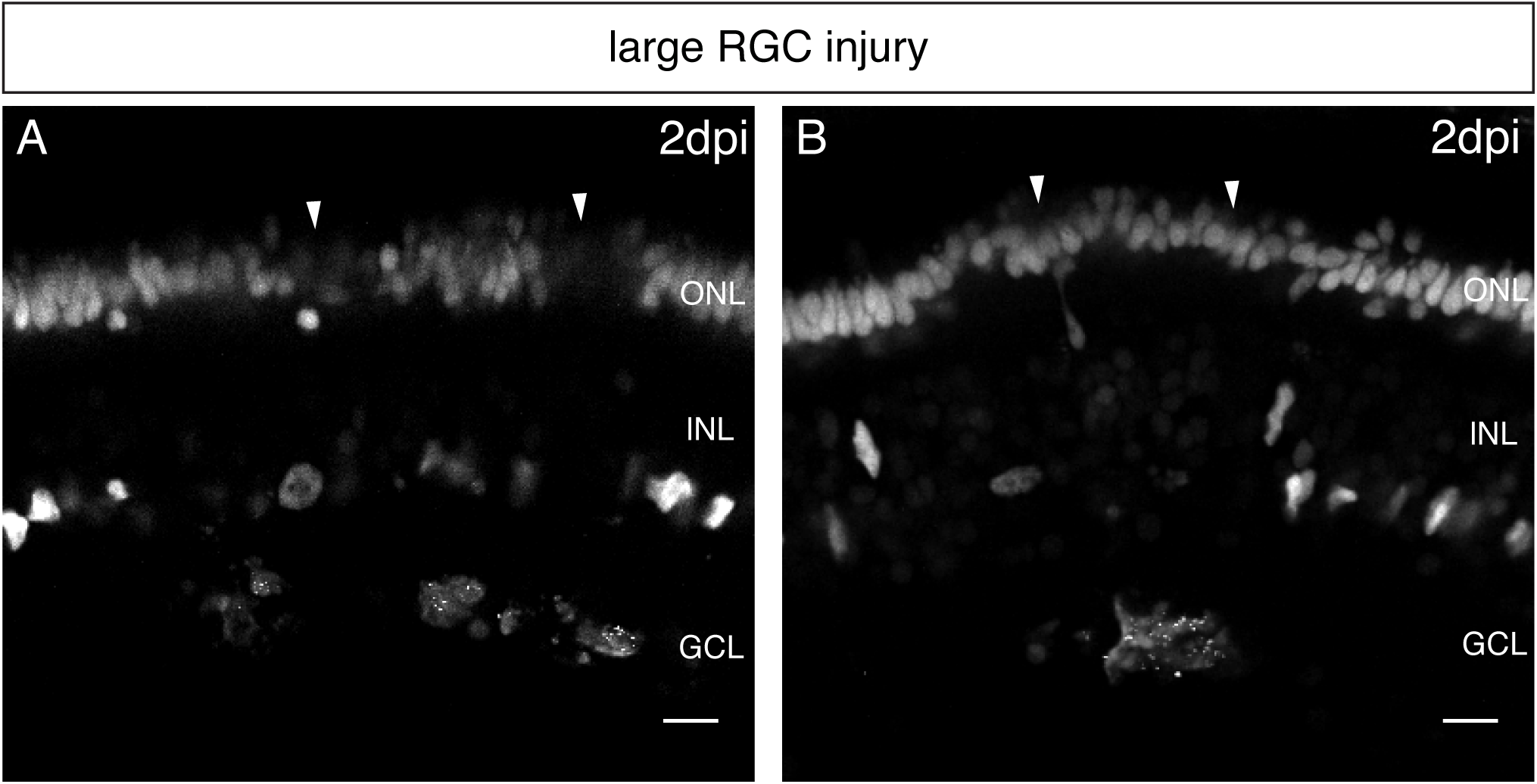
Increased RGC injuries lead to swelling and secondary cell death in the PRC layer. (A-B) *In vivo* imaging of juvenile *rx2*::H2B-eGFP medaka retinae which were either injured in the GCL (asterisks) using a two-photon laser and imaged 2 days later (n =10 fish, data obtained from five independent experiments). Large RGC injuries induce swelling and cell death in the PRC layer (arrowheads). olMG nuclei are largely depleted from the INL. Scale bars are 10 μm.

**Figure S5, related to Figure 5.**
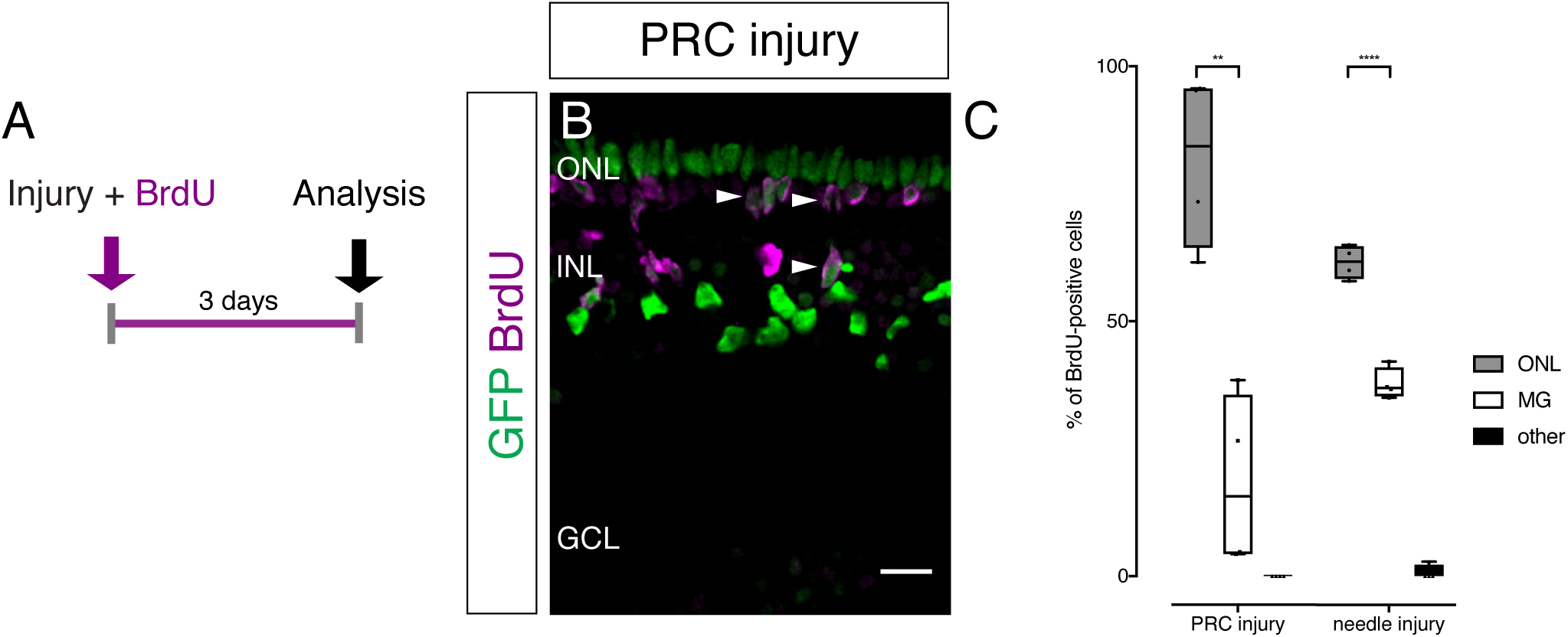
PRC and needle injuries trigger proliferation of olMG cells. (A) Scheme outlining the experimental procedure. Juvenile medaka were injured in the retina with a two-photon laser, ablating either PRCs or RGCs. The fish were incubated in BrdU for 3 days and analysed subsequently. (B) PRC injuries induce cell cycle re-entry of olMG cells detected by BrdU uptake. BrdU-positive nuclei are located in the INL (arrowheads) and below or in the ONL (arrowheads). Scale bar is 10 μm. (C) Counting of BrdU-positive nuclei and assigning them to different categories (ONL, olMG cells or other cell types) reveals individual profiles of the different injury types. BrdU-positive cells are mostly located in the ONL after PRC injuries (80%) and needle injuries (60%) whereas no BrdU-positive cells are detected after RGC injuries. **p=0.0019, ****p<0.0001. Box plots: median, 25th and 75th percentiles; whiskers show maximum and minimum data points. (n=3 fish each, data obtained from three independent experiments each).

**Figure S6, related to Figure 5.**
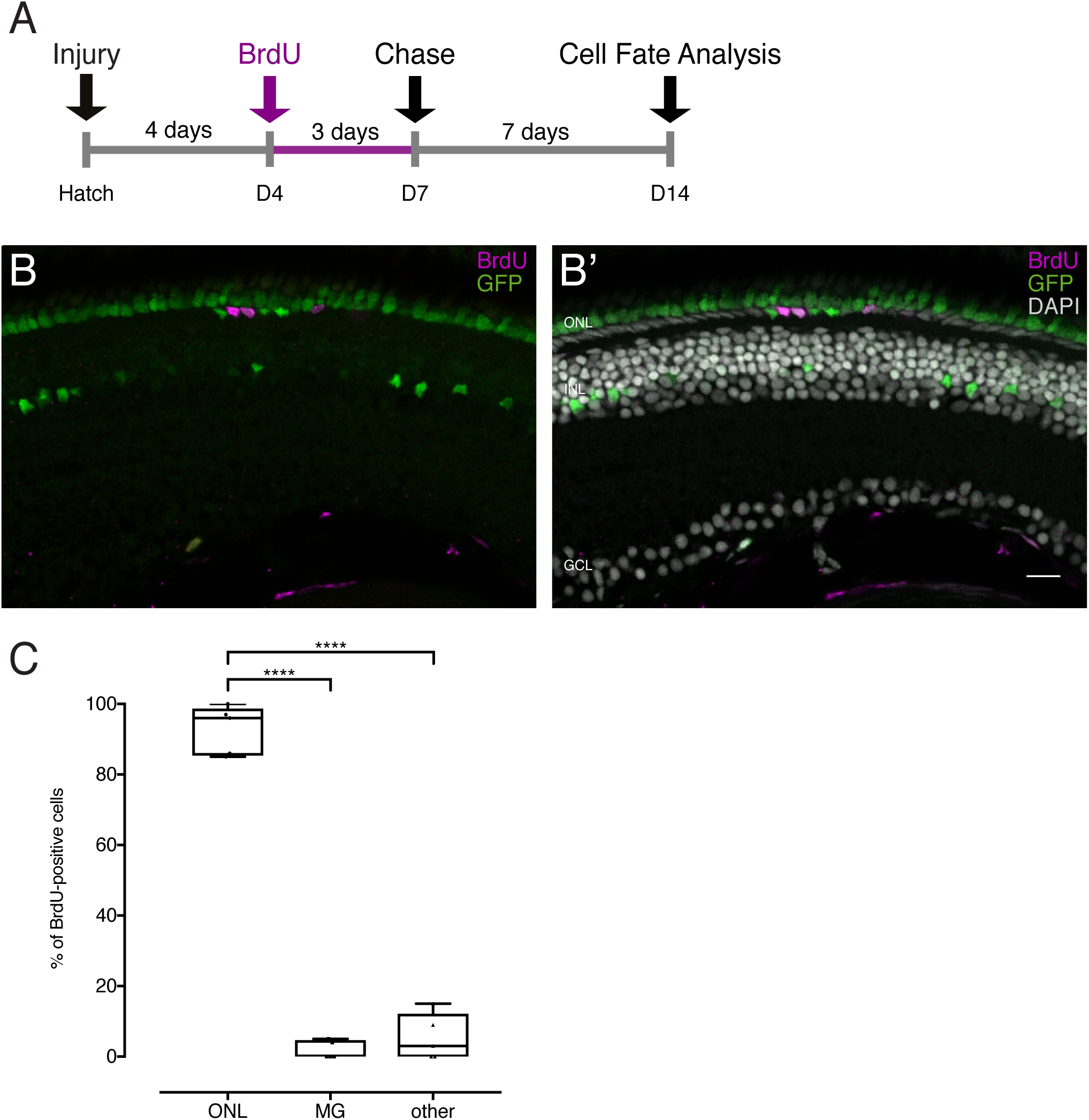
Late BrdU application after injury labels the same cell population as early BrdU application. (A) Scheme outlining the experimental procedure. Juvenile medaka were injured in the retina with a needle ablating all cell types. The fish were incubated in BrdU from 4 dpi until 7 dpi and analysed at 14 dpi. (B-C) BrdU-positive cells are located in the ONL, 1 out of 5 fish contained 1 BrdU-positive MG cell. GFP-positive olMG nuclei are depleted from the INL. ****p<0.0001. Box plots: median, 25th and 75th percentiles; whiskers show maximum and minimum data points (n=5 fish, data obtained from two independent experiments). Scale bar is 10 μm.

